# Structure of the Lifeact–F-actin complex

**DOI:** 10.1101/2020.02.16.951269

**Authors:** Alexander Belyy, Felipe Merino, Oleg Sitsel, Stefan Raunser

## Abstract

Lifeact is a short actin-binding peptide that is used to visualize filamentous actin (F-actin) structures in live eukaryotic cells using fluorescence microscopy. However, this popular probe has been shown to alter cellular morphology by affecting the structure of the cytoskeleton. The molecular basis for such artefacts is poorly understood. Here, we determined the high-resolution structure of the Lifeact–F-actin complex using electron cryo-microscopy. The structure reveals that Lifeact interacts with a hydrophobic binding pocket on F-actin and stretches over two adjacent actin subunits, stabilizing the DNase I-binding loop of actin in the closed conformation. Interestingly, the hydrophobic binding site is also used by actin-binding proteins, such as cofilin and myosin and actin-binding toxins, such as TccC3HVR from *Photorhabdus luminescens* and ExoY from *Pseudomonas aeruginosa*. *In vitro* binding assays and activity measurements demonstrate that Lifeact indeed competes with these proteins, providing an explanation for the altering effects of Lifeact on cell morphology *in vivo*. Finally, we demonstrate that the affinity of Lifeact to F-actin can be increased by introducing mutations into the peptide, laying the foundation for designing improved actin probes for live cell imaging.

## Introduction

The network of actin filaments in eukaryotic cells is involved in processes ranging from intracellular trafficking to cell movement, cell division and shape control (Pollard and Cooper, 2009). It is therefore not surprising that much effort has been directed to characterize the actin cytoskeleton under both physiological and pathological conditions. Numerous actin-visualizing compounds were developed to enable this. These include small molecules, labelled toxins, recombinant tags, as well as actin-binding proteins and peptides (see (Melak et al., 2017) for a detailed review). However, using these molecules to study actin *in vivo* often alters the properties of actin filaments to such an extent that normal homeostasis of the cytoskeleton is impaired. Since these side effects cannot be avoided, it is important to know their molecular basis in order to be able to adequately interpret the experimental data.

Phalloidin and jasplakinolide are cyclic peptides derived from the death cap mushroom *Amanita phalloides* and marine sponge *Jaspis johnstoni* (Crews et al., 1986; Lynen and Wieland, 1938), respectively. They bind specifically to F-actin, and when fused to a fluorescent probe their derivatives allow visualization of the cell cytoskeleton by fluorescence microscopy (Melak et al., 2017). However, both molecules strongly stabilize F-actin and shift the cellular actin equilibrium, largely limiting their use in live cell imaging (Bubb et al., 2000; Dancker et al., 1975). Recent cryo-EM studies from our group and others have uncovered how phalloidin and jasplakinolide affect the structure of F-actin and described the potential limitations of their use (Mentes et al., 2018; Merino et al., 2018a; Pospich et al., 2019).

The development of various fluorescent proteins provided new ways of visualizing the actin cytoskeleton. A simple and popular technique compatible with live cell imaging is to express actin fused to GFP-like proteins (Ballestrem et al., 1998). However, such actin chimeras often interfere with the normal functionality of the cytoskeleton in a way that results in experimental artefacts (Aizawa et al., 1997; Nagasaki et al., 2017). An alternative to GFP-actin is to fuse GFP to actin-binding proteins, such as utrophin (Burkel et al., 2007; Lin et al., 2011) or *Arabidopsis* fimbrin (Sheahan et al., 2004) and synthetic affimers that bind actin (Kost et al., 1998; Lopata et al., 2018). These actin filament markers have been successfully used in a variety of cell types and organisms (Melak et al., 2017; Montes-Rodriguez and Kost, 2017; Spracklen et al., 2014).

The most recent development is an F-actin binding nanobody called Actin-Chromobody that claims to have a minimal effect on actin dynamics and no notable effect on cell viability (Schiavon et al., 2019). However, the binding of the Actin-Chromobody to actin has not yet been characterized at molecular level, leaving the true extent of possible side effects open.

As described above, fluorophore-bound proteins are large and bulky resulting in possible steric clashes when interacting with actin. To avoid these problems, small fluorophore-labeled peptides were developed. The most commonly used one is Lifeact, which is a 17 amino acid peptide derived from the N-terminus of the yeast actin-binding protein ABP140. In the original publication, Lifeact was described as a novel F-actin probe that does not interfere with actin dynamics *in vitro* and *in vivo* (Riedl et al., 2008). The same group later reported transgenic Lifeact-GFP expressing mice that were phenotypically normal and fertile (Riedl et al., 2010), and no influence of Lifeact on cellular processes was found under the published experimental conditions. Two other groups performed a direct comparison of various F-actin binding probes and confirmed the low influence of Lifeact on cell cytoskeletal architecture (Belin et al., 2014; Sliogeryte et al., 2016). Later on, however, several major Lifeact-caused artefacts were described: Lifeact was unable to stain certain F-actin rich structures (Munsie et al., 2009; Sanders et al., 2013), it disturbed actin assembly in fission yeast (Courtemanche et al., 2016), it caused infertility and severe actin defects in *Drosophila* (Spracklen et al., 2014), and altered cell morphology in mammalian cells (Flores et al., 2019). The existing explanatory hypothesis suggests that Lifeact induces a conformational change in F-actin that affects binding of cofilin and eventually impairs cell cytoskeletal dynamics (Courtemanche et al., 2016). However, despite the widespread usage of Lifeact, the validity of this hypothesis is still a matter of debate since no structure of Lifeact-decorated F-actin has been available.

In order to address this, we solved the structure of the Lifeact–F-actin complex using single particle cryo-EM. The 3.5 Å structure reveals that Lifeact binds to the two consecutive actin subunits of the same strand of the filament and displaces the DNase I-binding loop (D-loop) upon binding. The binding site overlaps with that of cofilin and myosin, suggesting that artefacts in live-cell imaging are caused by competition between these proteins and Lifeact. Competition binding assays *in vitro* prove that this is indeed the case. Furthermore, we show that the binding of Lifeact to F-actin considerably reduces the *in vivo* toxicity of the actin-modifying toxin TccC3HVR from *Photorhabdus luminescens*. Our data will help to predict potential artefacts in experiments using Lifeact, and will serve as a strong basis for developing new actin-binding probes with improved properties.

## Results and Discussion

### Structure of the Lifeact–F-actin complex

Based on previous studies (Mentes et al., 2018; Merino et al., 2018b; Pospich et al., 2019), we know that phalloidin stabilizes actin filaments. When it is added during polymerization, the nucleotide binding pocket is occupied with an ADP and P_i_ and the D-loop is in the open conformation (Pospich et al., 2019). We therefore polymerized actin in the presence of phalloidin and added an excess of Lifeact to the formed filaments in order to fully decorate the filaments with Lifeact. We then determined the structure of this complex by cryo-EM (Fig. 1, Table 1). The average resolution of the reconstruction was 3.5 Å, with local areas reaching 3.0 Å (Fig. S1), which allowed us to build an atomic model in which we could position most of the side chains. We could clearly identify densities corresponding to ADP, Mg^2+^ and P_i_ in the nucleotide-binding pocket of actin (Fig. 1A) and a density corresponding to phalloidin at the expected position (Mentes et al., 2018; Merino et al., 2018a; Pospich et al., 2019) in the center of the filament (Fig. 1A). Lifeact was well resolved and we could unambiguously fit 16 out of 17 amino acids into the density (Fig. 1, Fig. S2).

**Figure 1.**
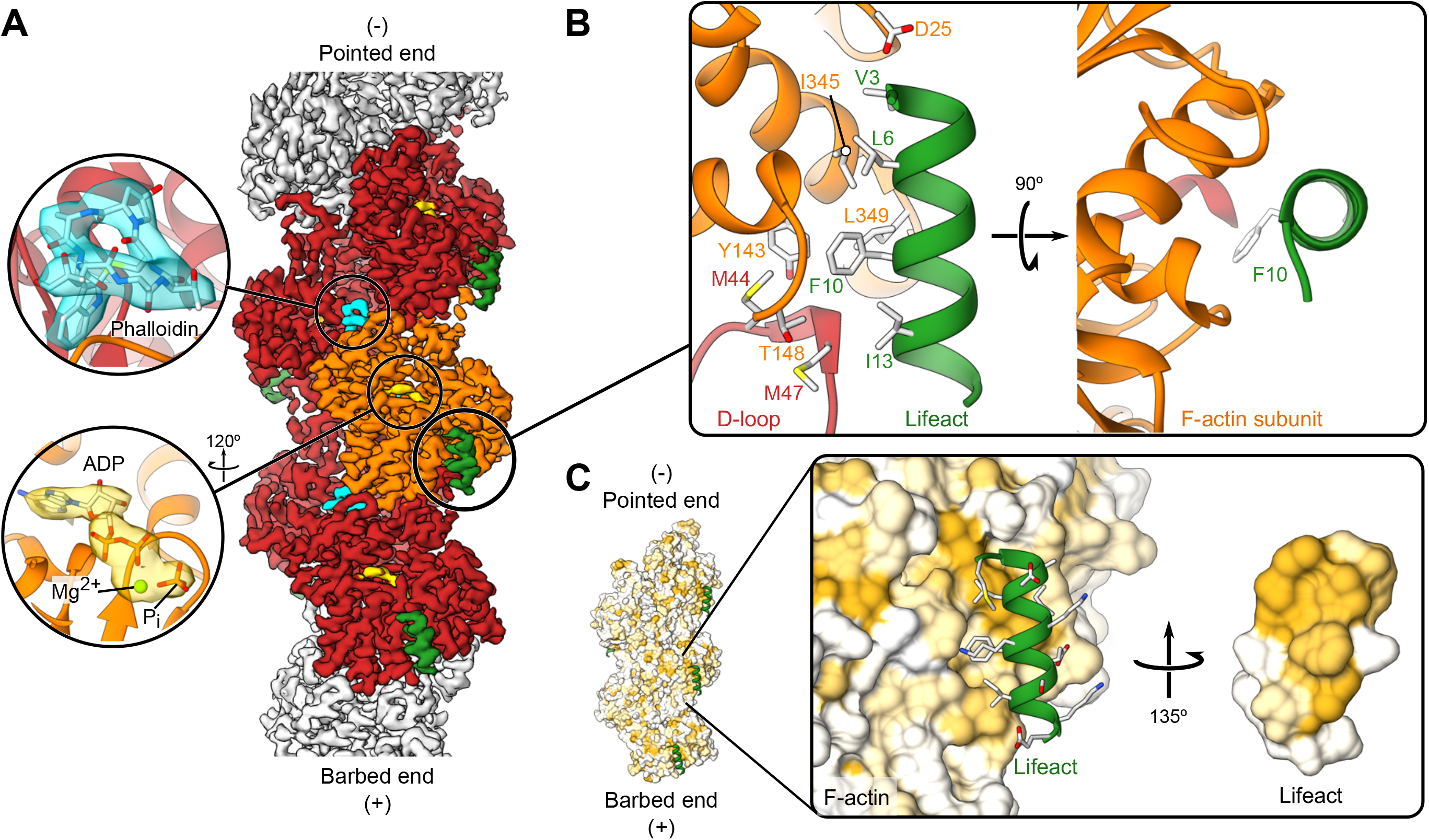
Cryo-EM structure of the Lifeact-F-actin-ADP-Pi-phalloidin complex. **A:** The 3.5 Å resolution map of the Lifeact-F-actin-ADP-P_i_-phalloidin complex shows a defined density for phalloidin (cyan), ADP-P_i_ (gold) and the Lifeact peptide (green). The central subunit of actin is colored in orange while its surrounding four neighbors are shown in red. **B**: Atomic model of the interface between Lifeact and F-actin. **C**: Surface of the atomic model of F-actin colored according to its hydrophobicity. Hydrophobicity increases as the color scale goes from white to gold. The inset highlights the hydrophobic nature of the Lifeact-binding surface.

**Table 1.**
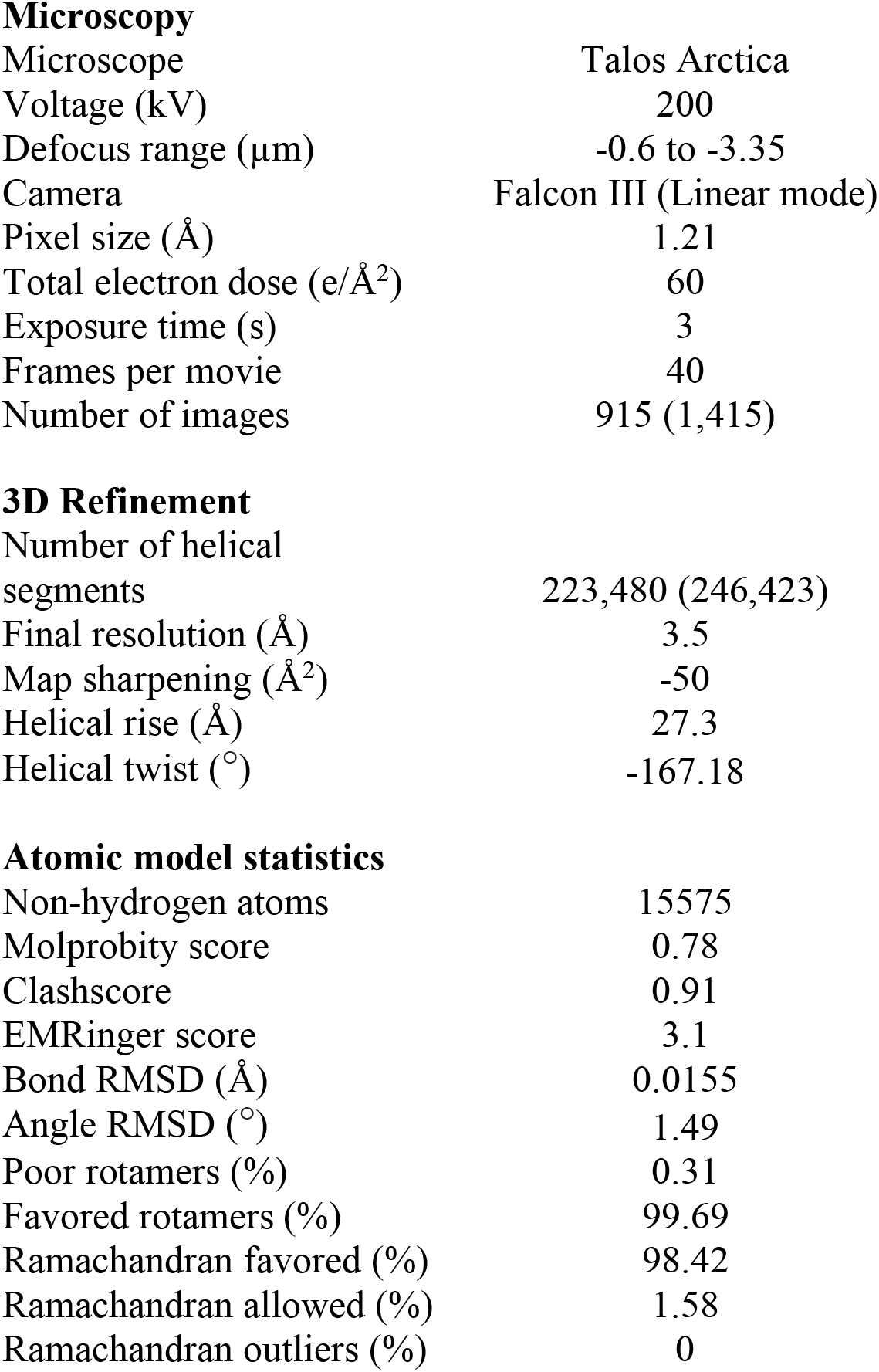
Cryo-EM data collection, refinement and validation statistics

Microscopy
|  |  |
| --- | --- |
| Microscope | Talos Arctica |
| Voltage (kV) | 200 |
| Defocus range ($\mu\text{m}$) | -0.6 to -3.35 |
| Camera | Falcon III (Linear mode) |
| Pixel size ($\text{\AA}$) | 1.21 |
| Total electron dose ($\text{e}/\text{\AA}^2$) | 60 |
| Exposure time (s) | 3 |
| Frames per movie | 40 |
| Number of images | 915 (1,415) |
| 3D Refinement |  |
| Number of helical segments | 223,480 (246,423) |
| Final resolution ($\text{\AA}$) | 3.5 |
| Map sharpening ($\text{\AA}^2$) | -50 |
| Helical rise ($\text{\AA}$) | 27.3 |
| Helical twist ($^\circ$) | -167.18 |
| Atomic model statistics |  |
| Non-hydrogen atoms | 15575 |
| Molprobity score | 0.78 |
| Clashscore | 0.91 |
| EMRinger score | 3.1 |
| Bond RMSD ($\text{\AA}$) | 0.0155 |
| Angle RMSD ($^\circ$) | 1.49 |
| Poor rotamers (%) | 0.31 |
| Favored rotamers (%) | 99.69 |
| Ramachandran favored (%) | 98.42 |
| Ramachandran allowed (%) | 1.58 |
| Ramachandran outliers (%) | 0 |

The peptide folds as an α-helix and spans two consecutive actin subunits of the same strand of the filament (Fig. 1A, B). The binding pocket is formed by the tip of the D-loop of the lower subunit (M47) and SD1 of the upper subunit, where the N-terminal region of Lifeact is almost locked in by the protruding D25 of actin (Fig. 1B). Although Lifeact is in general a hydrophilic peptide, it contains a hydrophobic patch formed by the side chains of V3, L6, I7, F10 and I13 which all orient to one side. This hydrophobic patch interacts with a hydrophobic groove on the surface of F-actin which comprises M44, M47, Y143, I345 and L349. F10 of Lifeact is deeply buried in this pocket (Fig. 1C). Interestingly and contrary to what we have seen before in samples co-polymerized with phalloidin (Pospich et al., 2019), the D-loop is in its closed conformation (Fig. S3). A comparison between the Lifeact-F-actin-ADP-P_i_ - phalloidin-structure with that of phalloidin-stabilized F-actin-ADP-P_i_ (Pospich et al., 2019) shows that direct interactions between Lifeact and the D-loop of F-actin are only possible if the D-loop is in its closed conformation (Movie S1). Specifically, I13 of Lifeact interacts with M47 of F-actin, stabilizing the closed D-loop conformation in F-actin. This suggests that Lifeact has a higher affinity to F-actin-ADP, where the D-loop is in the closed conformation, than to phalloidin-stabilized F-actin-ADP-P_i_, where the D-loop has to be first moved from the open to the closed conformation. Indeed, Kumari *et al*. (Kumari et al., 2019) showed in a complementary study that the affinity of Lifeact is three to four times higher for F-actin-ADP compared to F-actin-ADP-P_i_.

### Properties of the interaction site

Guided by the insights gained from our structure, we mutated different residues at the peptide-actin interface to study the binding properties of Lifeact in more detail. We chose to use *Saccharomyces cerevisiae* for these studies since actin mutagenesis can be easily and rapidly performed in this organism. To avoid toxicity from artificial overexpression of Lifeact, we expressed Lifeact-mCherry under the promoter of the actin binding protein ABP140 from which Lifeact was originally derived, and then performed confocal microscopy experiments to visualize Lifeact-actin interaction.

When expressing WT Lifeact-mCherry, we observed the typical patch morphology of actin-rich structures that are distinct from the diffuse background of Lifeact-mCherry. These actin-rich patches can also be observed when the cells are stained by fluorescently-labelled phalloidin (Fig. 2A). However, when L6 was mutated to lysine, or F10 to alanine, we did not observe these structures and Lifeact-mCherry was homogeneously distributed in the cells. Although the I13A variant displayed some of the patches, they were significantly less abundant than with WT Lifeact-mCherry (Fig. 2A, B). While the Lifeact L6K mutant introduces a charge in the hydrophobic patch, Lifeact mutants F10A and I13A retain the hydrophobicity but change the surface structure of the peptide. Since all mutations impaired the interaction between actin and Lifeact, we conclude that hydrophobicity as well as shape complementarity are important for efficient Lifeact binding to F-actin.

**Figure 2.**
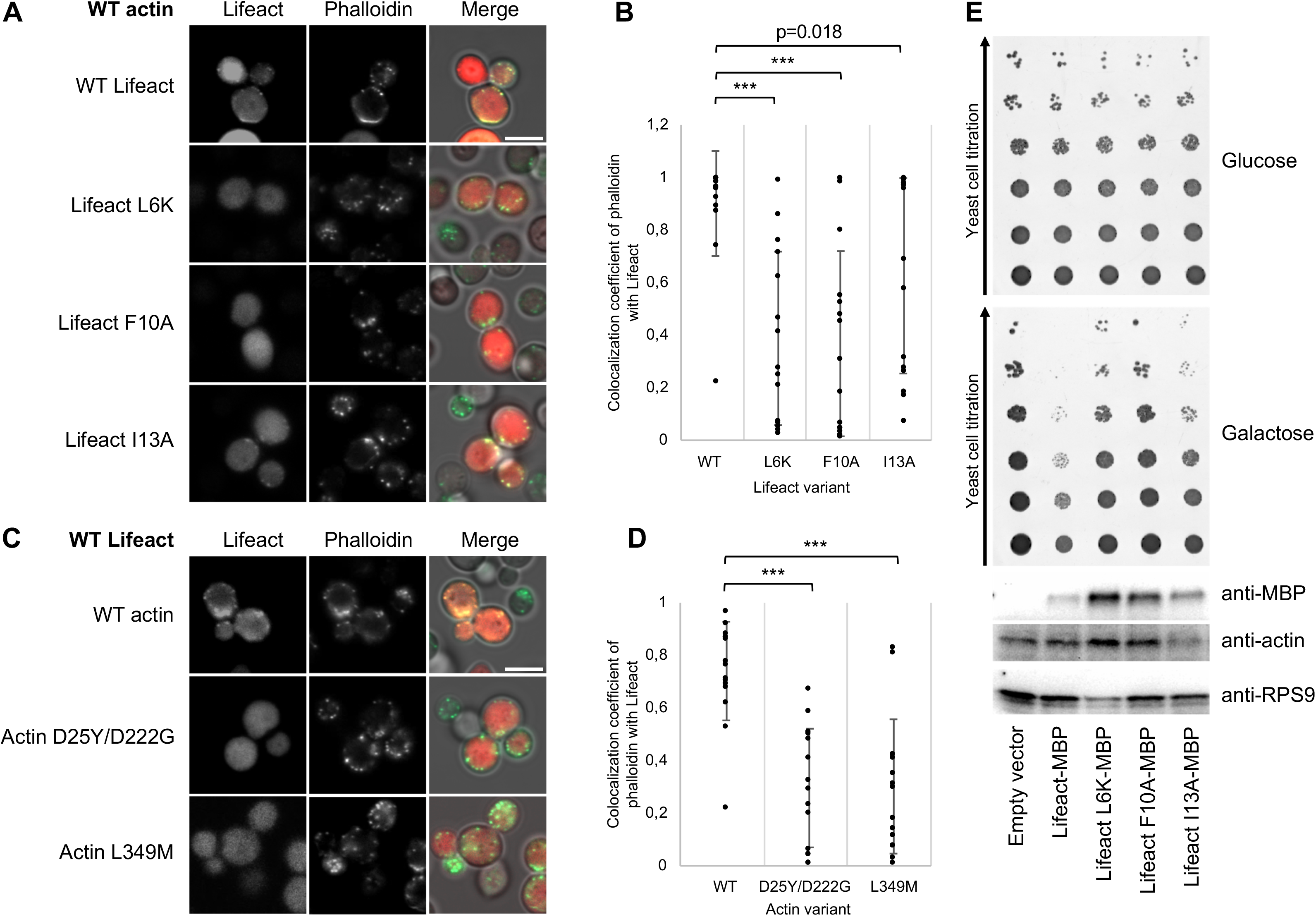
The Lifeact–F-actin complex is affected by point mutations. **A, C**: Confocal microscopy images of yeast cells expressing Lifeact-mCherry variants in a WT actin background (**A**) and WT Lifeact-mCherry in yeast with different actin variants (**C**). Actin was additionally stained with fluorescently labeled phalloidin (ActinGreen 488). Representative areas where yeast cells are stained with both fluorophores are shown. Note that for our experiments we used the previously described D25Y/D222G double mutant of yeast actin (Belyy et al., 2016). However, D222 is located in subdomain IV and is therefore unlikely to play a role in the Lifeact–F-actin interaction. Scale bars, 5 μm. **B, D**: Calculated weighted colocalization coefficients of phalloidin with Lifeact-mCherry from 15 yeast cells from two independent experiments with five micrographs each, corresponding to (**A**) and (**C**), respectively. For statistical analysis, the paired *t* test was used. *** p < 0.001. The error bars in the panels correspond to standard deviations of three independent experiments. **E**: Growth phenotype assay with yeast overexpressing Lifeact-MBP variants under a strong galactose promoter. The top image marked “Glucose” corresponds to experimental conditions with low Lifeact expression. The central image marked “Galactose” corresponds to experimental conditions with high Lifeact expression. The lower image is a Western blot of cells grown on galactose-containing media performed using anti-MBP, anti-actin and anti-RPS9 antibodies.

Our structure suggests that actin D25 acts as an N-terminal cap for the helix of Lifeact and a mutation of actin D25 to tyrosine would affect this interaction and mutating L349 of actin to methionine would impair its crucial interaction with Lifeact F10. Indeed, actin-rich structures were also absent when Lifeact-mCherry was expressed in cells with the D25Y or the L349M actin variant (Fig. 2C, D), indicating that actin D25 and Lifeact F10 are important for Lifeact binding.

To study the effect of Lifeact WT and variants on yeast viability we overexpressed Lifeact-MBP fusions under a strong galactose promoter and analyzed their toxicity in a yeast growth phenotype assay. Consistent with a previously reported study (Courtemanche et al., 2016), we observed that the overexpression of Lifeact-MBP caused cell toxicity (Fig. 2E). However, mutagenesis of I13 to alanine improved, and L6 to lysine and F10 to alanine fully restored yeast growth. Altogether, these results demonstrate the importance of shape complementarity as well as hydrophobicity at the Lifeact-actin interface.

### Lifeact mutations increase its affinity to F-actin

Despite these specific interactions between Lifeact and F-actin, the peptide binds to F-actin only with micromolar affinity (Riedl et al., 2008). A higher affinity would provide a stronger signal-to-noise ratio, decreasing the background during live imaging, and allowing lower expression levels of the peptide to be used during such experiments. We therefore attempted to increase the affinity of Lifeact to F-actin by structure-guided *in silico* design using RosettaScripts (Fleishman et al., 2011) based on our atomic model. The simulation output suggested several possible mutations after residue 12 of Lifeact (Fig. 3A). The mutation E16R was especially promising. It was predicted to add an additional interaction with the D-loop and an electrostatic interaction with E167 of actin (Fig. 3B). Indeed, this variant showed an increased affinity for F-actin as judged by cosedimentation assays (Fig. 3C, D). Although we could not observe density for E17 of Lifeact in our density map, we also created and tested a E17K variant of Lifeact which should similarly create an additional interaction with E167 of actin and thereby increase the affinity of the peptide. In line with the prediction, E17K Lifeact-MBP variants showed an increased affinity for F-actin (Fig. 3C, D). Together, these modifications show that based on our atomic model, Lifeact can be optimized by mutations to increase its binding to actin.

**Figure 3.**
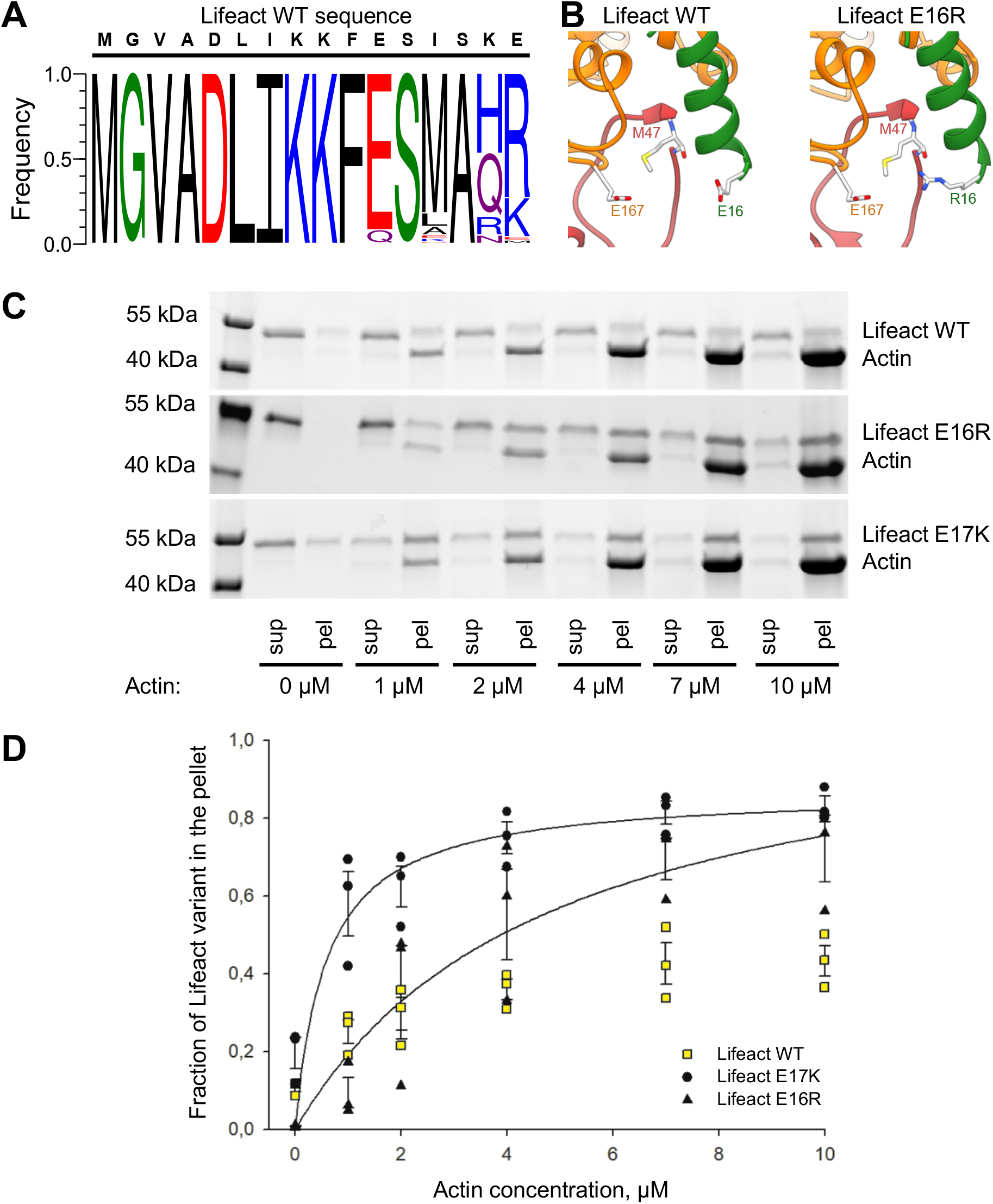
Lifeact sequence design. **A**: Frequency of amino acids in the top 100 designs produced by Rosetta. **B**: Predicted structure of the E16R mutant. The wild-type structure is included for comparison. **C**: Cosedimentation of F-actin and 1 μM Lifeact-MBP proteins detected by SDS-PAGE. The upper band corresponds to Lifeact-MBP, and the lower band corresponds to actin. sup, supernatant; pel, pellet. A representative stain-free gel is shown. **D**: The fractions of Lifeact-MBP that cosedimented with F-actin were quantified by densitometry and plotted versus actin concentrations. The error bars in panel **D** correspond to standard deviations of three independent experiments.

### Lifeact competes with cofilin and myosin

Several studies demonstrated that Lifeact staining interfered with cofilin binding to actin. It was shown that Lifeact does not bind to cofilin-bound F-actin in cells (Munsie et al., 2009) and Lifeact-expressing cells possess longer and thicker stress fibers. Studies by Flores *et al*. (Flores et al., 2019) suggested that reduced cofilin binding to F-actin is the underlying cause of the observed Lifeact-induced artefacts. Similarly, Lifeact caused changes in endocytosis and cytokinesis of *Schizosaccharomyces pombe*, which were attributed to reduced cofilin interaction with actin (Courtemanche et al., 2016). The authors of that study proposed that cofilin and Lifeact bind to different regions of F-actin, and suggested that binding of one of these proteins impairs binding of the other by provoking a conformational change in F-actin (Courtemanche et al., 2016).

Apart from Lifeact-induced stabilization of the closed conformation of the D-loop, however, our structure does not show major differences to previously reported structures of F-actin (Merino et al., 2018b; Pospich et al., 2019). Therefore, a conformational change in F-actin cannot be the cause for the effect of Lifeact. When comparing our F-actin–Lifeact structure with that of F-actin–cofilin (Tanaka et al., 2018), it becomes obvious that the binding site of cofilin overlaps with that of Lifeact (Fig. 4A). Notably, the same is true for myosin, which interacts with the same position on the actin surface (Fig. 4B) (Ecken et al., 2016). We therefore performed *in vitro* competition actin binding assays with human cofilin-1, the motor domain of human non-muscle myosin 2C isoform (NM2C), and Lifeact. Lifeact successfully decreased cofilin and myosin binding in a dose-dependent manner (Fig. 4C-F). As a negative control, we performed a similar competition assay with tropomyosin that binds to a different region of actin (Fig. 4G) (Ecken et al., 2015), and the Lifeact F10A mutant which only binds weakly to F-actin (Fig. 4C-F). As expected, the addition of Lifeact did not affect tropomyosin binding to F-actin (Fig. 4H, I), nor could F10A Lifeact compete with cofilin-1 or NM2C (Fig. 4C-F). Based on our structural and functional data, we demonstrate that the morphological artefacts described for Lifeact are not due to a conformational change in actin but are caused by competition for the same binding site on F-actin of Lifeact with actin-binding proteins, such as cofilin and myosin.

**Figure 4.**
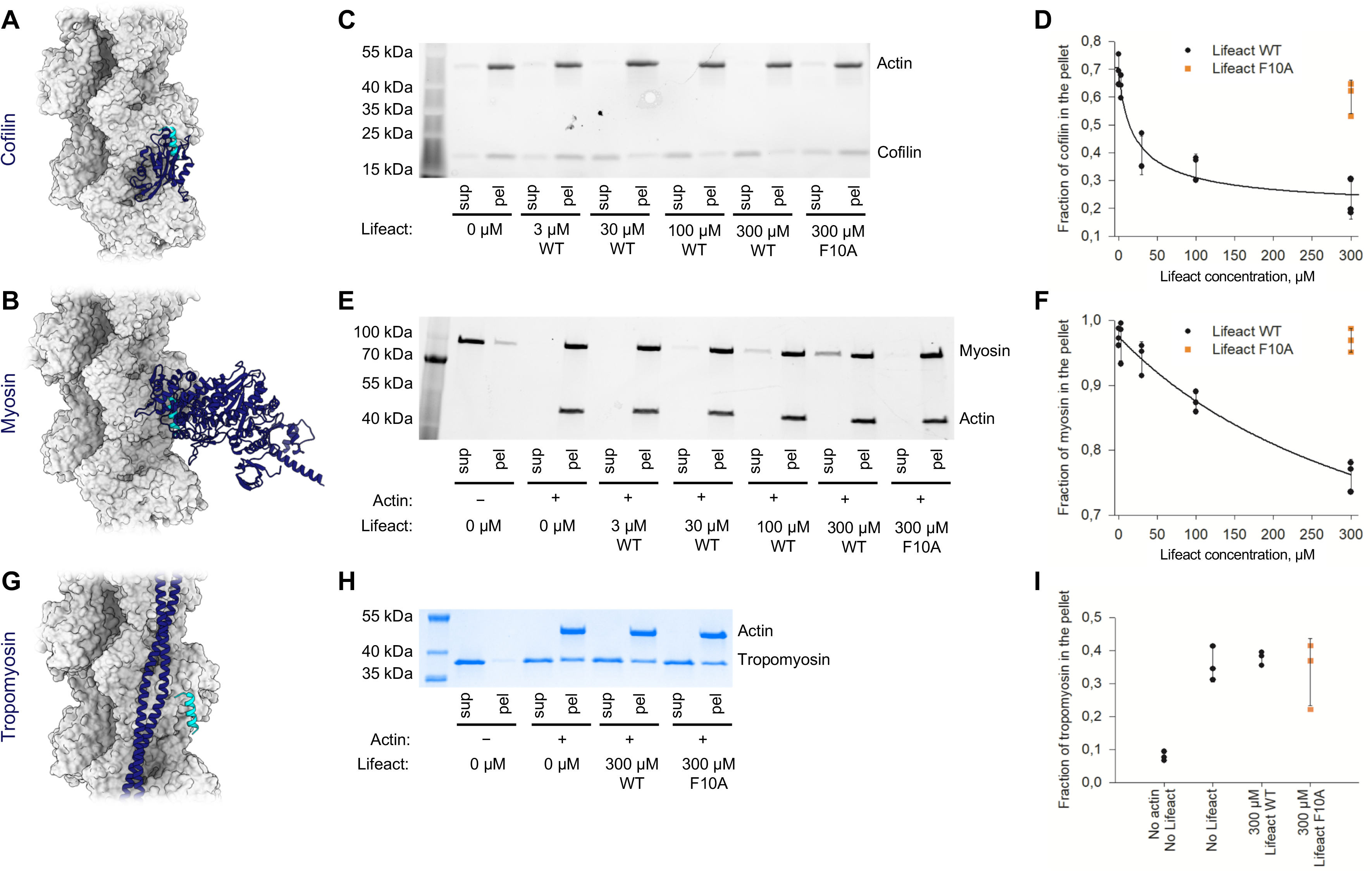
Lifeact competes with cofilin and myosin *in vitro*. **A, B, G**: Structural models of the (**A**) cofilin–F-actin (PDB 5YU8) (Tanaka et al., 2018), (**B**) myosin–F-actin (PDB 5JLH) (Ecken et al., 2016) and (**G**) tropomyosin–F-actin (3J8A) (Ecken et al., 2015) complexes. **C**: SDS-PAGE analysis of cosedimentation experiments of F-actin (3 μM, upper band) with cofilin (3 μM, lower band) in the presence of the indicated amounts of Lifeact. A representative stain-free gel is shown. **E**: SDS-PAGE analysis of cosedimentation experiments of F-actin (0.7 μM, lower band) with myosin (0.5 μM, upper band) in the presence of the indicated amounts of Lifeact. A representative stain-free gel is shown. **H**: SDS-PAGE analysis of cosedimentation experiments of F-actin (3 μM, lower band) with tropomyosin (3 μM, upper band) in the presence of the indicated amounts of Lifeact. A representative Coomassie stained gel is shown. The fractions of cofilin, myosin and tropomyosin that co-sedimented with F-actin in the corresponding experiments were quantified by densitometry and plotted against Lifeact concentrations at **D**, **F** and **I**, respectively. The error bars in **D**, **F** and **I** correspond to standard deviations of three independent experiments. sup, supernatant; pel, pellet.

### Lifeact impairs the activity of bacterial toxins

In our laboratory, we study two bacterial toxins that interact with F-actin. One is *Pseudomonas aeruginosa* ExoY, a toxin that becomes a potent nucleotidyl cyclase upon interaction with Factin (Belyy et al., 2016). After activation, the toxin generates a supraphysiologic amount of cGMP and cAMP that impedes cell signaling. It was previously demonstrated that the mutagenesis of D25 in actin abolishes ExoY binding to F-actin (Belyy et al., 2018). The same actin mutation also prevents Lifeact binding, we therefore hypothesize that Lifeact and ExoY have overlapping binding sites.

The second toxin is the 30 kDa C-terminal fragment of *Photorhabdus luminescens* TccC3 (TccC3HVR), which is the effector domain of the large Tc toxin complex PTC3. Once it is translocated into the cell by the injection machinery of PTC3, TccC3HVR acts as an ADP-ribosyltransferase that modifies actin at T148 (Lang et al., 2010). This leads to uncontrolled actin polymerization, clustering, and finally to cell death due to cytoskeletal collapse. T148 is located in close proximity to the Lifeact binding site, therefore the actin binding site of TccC3HVR and Lifeact might overlap.

To understand whether Lifeact competes with the binding of ExoY, we first performed a cosedimentation assay with ExoY, F-actin and different concentrations of Lifeact. In agreement with our hypothesis, we observed a decrease of ExoY binding to F-actin in the presence of Lifeact while the Lifeact F10A mutant did not impair formation of the ExoY-F-actin complex (Fig. 5A, B).

**Figure 5.**
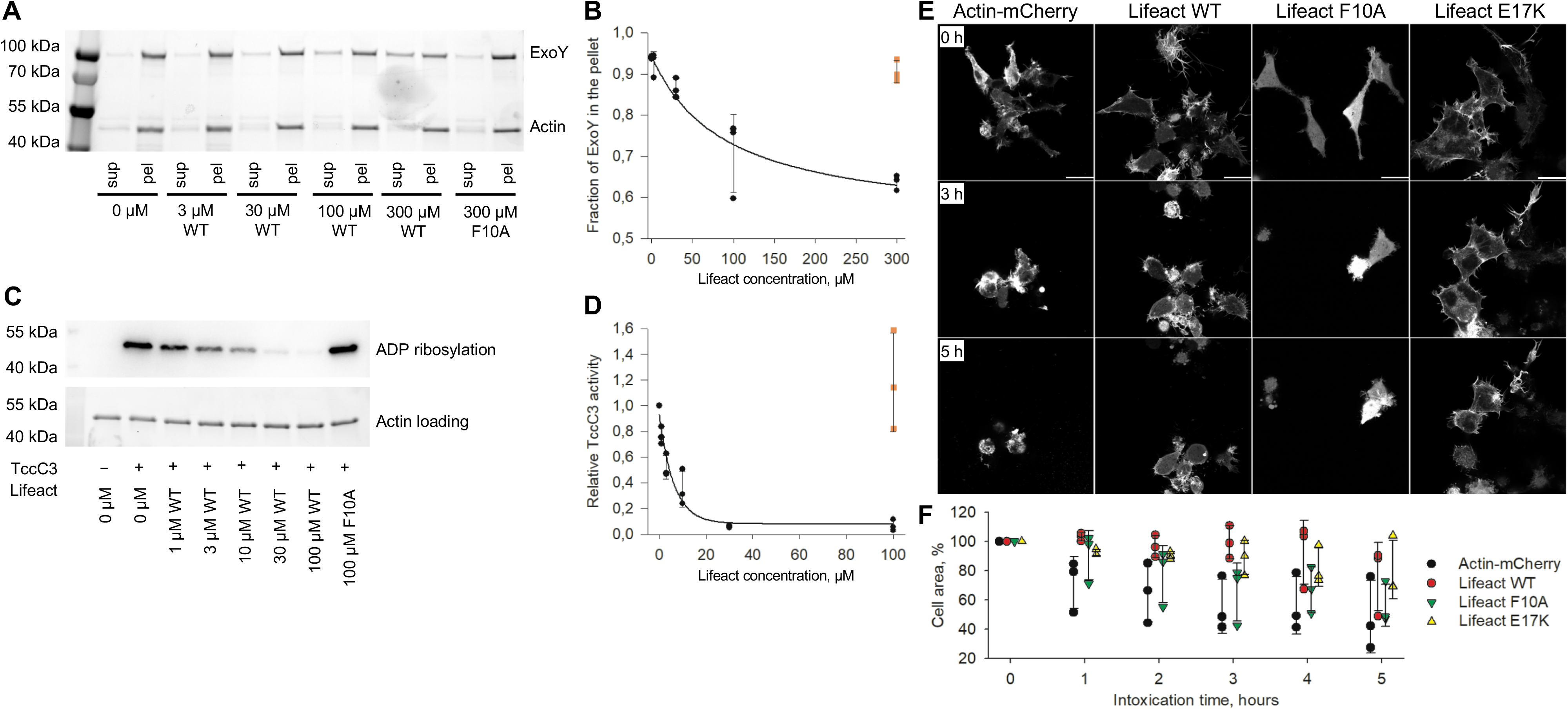
Lifeact impairs the activity of F-actin binding bacterial toxins. **A**: SDS-PAGE analysis of cosedimentation experiments of F-actin (1 μM, lower band) with ExoY-MBP (1 μM, upper band) in the presence of the indicated amounts of Lifeact. sup, supernatant; pel, pellet. A representative stain-free gel is shown. The fractions of ExoY that co-sedimented with F-actin were quantified by densitometry and plotted against Lifeact concentrations in **B**. **C**: Level of actin ADP-ribosylation by TccC3HVR in the presence of Lifeact was analyzed by Western blot using an ADP-ribose binding reagent. The equal loading of actin was additionally checked by imaging the same stain-free gel prior to blotting (lower image). The ADP-ribosylation level of actin was quantified by densitometry and plotted against Lifeact concentrations in **D**. Error bars at **B** and **D** correspond to standard deviations of three independent experiments. **E**: HEK 293T cells expressing mCherry fusions of actin or LifeAct variants were intoxicated with 300 pM of the *P. luminescens* toxin PTC3, which injects TccC3HVR into cells. The degree of cytoskeletal collapse and accompanying cell shrinkage was monitored for 5 h using live cell imaging, and is plotted in **F** based on three independent experiments for each condition. Scale bars, 20 μm.

We then ADP-ribosylated F-actin by TccC3HVR in the presence of Lifeact. In our experimental setup, 3 μM of WT Lifeact was already sufficient to decrease the level of ADP ribosylation by a factor of two, while in the control reaction 100 μM of F10A Lifeact did not decrease the level of ADP-ribosylation at all (Fig. 5C, D). This experiment strongly supports the hypothesis that TccC3HVR and Lifeact bind to the same region of F-actin.

Encouraged by these *in vitro* results, we decided to test whether expressing Lifeact in mammalian cells would protect them from the TccC3HVR toxin. We therefore expressed either mCherry-tagged WT Lifeact, Lifeact F10A, Lifeact E17K, or mCherry-tagged actin as a negative control in adherent HEK 293T cells. We then intoxicated the cells with PTC3 and observed the effect of the injected TccC3HVR. Our control cells expressing actin and cells expressing the F-actin binding-incompetent F10A Lifeact showed rapid cytoskeletal collapse and accompanying overall shrinkage (Fig. 5E, F). However, the toxic effect of TccC3HVR was significantly reduced in cells that expressed WT Lifeact or the binding-competent E17K mutant. Thus, Lifeact has anti-toxin properties and despite its effects on the cytoskeleton, it has the potential to be used as a precursor for the development of anti-toxin drugs.

## Conclusions

In this study we determined the binding site of Lifeact on F-actin and demonstrated that this peptide directly competes with actin-binding proteins such as cofilin and myosin, providing an explanation for how Lifeact alters cell morphology. In addition, we demonstrate how the affinity of Lifeact can be modulated by site directed mutagenesis in order to create Lifeact-based probes with modified properties. Our results have strong implications for the usage of Lifeact as an actin filament label in fluorescent light microscopy and provide cell biologists with the background information that is needed to make a properly informed decision on whether to use Lifeact in an experiment. Furthermore, we have demonstrated that Lifeact competes with actin-binding toxins such as ExoY and TccC3HVR, and partially counteracts the intoxication of cells by PTC3 toxin. This paves the way for the development of Lifeact-based anti-toxin drugs.

## Materials and methods

### Plasmids, bacteria and yeast strains, growth conditions

The complete list of used oligonucleotides, constructions and strains can be found in the Supplementary data. *E. coli* strains were grown in LB medium supplemented with ampicillin (100 μg/ml) or kanamycin (50 μg/ml). *S. cerevisiae* were grown on rich YPD medium or on synthetic defined medium (Yeast nitrogen base, Difco) containing galactose or glucose and supplemented if required with uracil, histidine, leucine, tryptophan, or adenine. *S. cerevisiae* strains were transformed using the lithium-acetate method (Daniel Gietz and Woods, 2002). Yeast actin mutagenesis was performed as described previously (Belyy et al., 2015). Yeast viability upon Lifeact-MBP overexpression under the galactose promoter was analyzed by a drop test: 5-fold serial dilutions of cell suspensions were prepared from overnight agar cultures by normalizing OD_600_ measurements, then spotted onto agar plates and incubated for 2-3 days at 30 °C. Analysis of protein expression in yeast was performed following the described protocol (Kushnirov, 2000): yeast cells were grown in liquid galactose-containing medium overnight at 30 °C. Cells corresponding to 1 ml of OD_600_ 1.0 were washed with 0.1 M NaOH, resuspended in 50 μl of 4-fold Laemmli sample buffer, and boiled for 5 minutes at 95 °C. 5 μl of the extracts were separated by SDS-PAGE, followed by Western blotting analysis and incubation with anti-MBP (NEB), anti-actin (C4, Abcam) or anti-RPS9 serum (polyclonal rabbit antibodies were a generous gift of Prof. S. Rospert).

### Protein expression and purification

Fusion proteins of Lifeact variants or *Pseudomonas aerugino*sa ExoY toxin and maltose-binding protein (MBP) were purified from *E. coli* BL21-CodonPlus(DE3)-RIPL cells harboring the corresponding plasmids (2489 pB502 WT Lifeact-MBP, 2490 pB506 E16R Lifeact-MBP, 2491 pB524 E17K Lifeact-MBP, 2479 pB386 MBP-ExoY). A single colony was inoculated in 100 ml of LB media and grown at 37 °C. At OD_600_ 1.0 protein expression was induced by addition of IPTG to a final concentration of 1 mM. After 2 h of expression at 37 °C, the cells were harvested by centrifugation, resuspended in buffer A (20 mM Tris pH 8, 500 mM NaCl), and lysed by sonication. The soluble fraction was applied on buffer A-equilibrated Protino Ni-IDA resin (Macherey-Nagel), washed, and eluted by buffer A, supplemented with 250 mM imidazole. Finally, the eluates were dialyzed against buffer B (20 mM Tris pH 8, 150 mM NaCl) and stored at −20 °C.

Rabbit skeletal muscle α-actin was purified as described previously (Merino et al., 2018b) and stored in small aliquots at −80 °C.

Human cofilin-1 was purified from *E. coli* cells using previously described method (Carlier et al., 1997). In short, Rosetta DE3 *E. coli* cells were transformed with the 1855 plasmid. An overnight culture derived from a single colony was diluted into 2 L of LB media to OD_600_ 0.06 and grown at 37 °C. When OD_600_ reached 0.7, the cells were cooled to 30 °C and cofilin expression was induced by adding IPTG to a final concentration of 0.5 mM. After 4 h of expression, the cells were harvested by centrifugation, resuspended in buffer C (10 mM Tris pH 7.8, 1 mM EDTA, 1 mM PMSF and 1 mM DTT), and lysed using a fluidizer. The soluble fraction of the lysate was dialyzed overnight in buffer D (10 mM Tris pH 7.8, 50 mM NaCl, 0.2 mM EDTA and 2 mM DTT), and cleared by centrifugation. Then, the lysate was applied onto DEAE resin and washed with buffer D. Cofilin-containing fractions of the flow-through were collected and dialyzed against buffer E (10 mM PIPES pH 6.5, 15 mM NaCl, 2 mM DTT and 0.2 mM EDTA). After centrifugation, the protein was loaded onto Mono S cation exchange column and eluted by a linear gradient of 15 mM to 1 M NaCl in buffer E. Cofilin-containing fractions were concentrated to 10 mg/ml and stored at −80 °C.

Human tropomyosin was purified from *E. coli* BL21(DE3) cells transformed with a 1609 plasmid using the previously described method (Coulton et al., 2006) with minor modifications. In brief, an overnight culture derived from a single colony was diluted into 5 L of LB media to OD_600_ 0.06 and grown at 37 °C. When the OD_600_ reached 0.5, the cells were cooled down to 20 °C and recombinant protein expression was induced by adding IPTG to a final concentration of 0.4 mM. After overnight protein expression, the cells were harvested by centrifugation, resuspended in buffer F (20 mM Tris pH 7.5, 100 mM NaCl, 5 mM MgCl_2_, 2 mM EGTA and Roche cOmplete protease inhibitor) and lysed by fluidizer. The soluble fraction of the lysate was heated for 10 minutes at 80 °C, then cooled down to 4 °C and centrifuged. The supernatant was mixed 1:1 with buffer H (20 mM sodium acetate buffer pH 4.5, 100 mM NaCl, 5 mM MgCl_2_ and 2 mM EGTA). The precipitate was collected and incubated for 1 h with buffer I (10 mM Bis-Tris pH 7, 100 mM NaCl). The renatured protein was applied to a HiTrap Q anion exchange column and eluted by a linear gradient of 100 mM to 1 M NaCl in buffer I. Tropomyosin-containing fractions were pooled and stored at −80 °C.

The motor domain of non-muscular myosin-2C (MYH14, isoform 2 from *H. sapiens*) consisting of amino acids 1–799 was purified as described previously (Ecken et al., 2016) Tcc3HVR, the ADP-ribosyltransferase domain of the *Photorhabdus luminescens* TccC3 protein (amino acids 679 - 960) was purified as described previously (Roderer et al., 2019).

The TcdA1 and TcdB1-TccC3 components of the *Photorhabdus luminescens* PTC3 toxin were expressed and purified as described previously (Gatsogiannis et al., 2016).

### Cryo-EM sample preparation, data acquisition, and processing

Actin was polymerized by incubation in F-buffer in the presence of a twofold molar excess of phalloidin for 30 minutes at room temperature and further overnight at 4 °C. The next day, actin filaments were pelleted using a TLA-55 rotor for 30 minutes at 150.000 g at 4 °C and resuspended in F-buffer. 5 minutes before plunging, F-actin was diluted to 6 μM and mixed with 200 μM Lifeact peptide (the peptide of sequence MGVADLIKKAESISKEE with C-terminal amide modification was provided by Genosphere with >95% purity. The peptide was dissolved in 10 mM Tris pH 8). To improve ice quality, Tween-20 was added to the sample to a final concentration of 0.02% (w/v). Plunging was performed using the Vitrobot Mark IV system (Thermo Fisher Scientific) at 13 °C and 100% humidity: 3 μl of sample were applied onto a freshly glow-discharged copper R2/1 300 mesh grid (Quantifoil), blotted for 8 s on both sides with blotting force −20 and plunge-frozen in liquid ethane.

The dataset was collected using a Talos Arctica transmission electron microscope (Thermo Fisher Scientific) equipped with an XFEG at 200 kV using the automated data-collection software EPU (Thermo Fisher Scientific). Two images per hole with defocus range of −0.6 - −3.35 μm were collected with the Falcon III detector (Thermo Fisher Scientific) operated in linear mode. Image stacks with 40 frames were collected with total exposure time of 3 sec and total dose of 60 e^−^/Å. 1415 images were acquired and 915 of them were further processed. Filaments were automatically selected using crYOLO (Wagner et al., 2019). Classification, refinement and local resolution estimation were performed in SPHIRE (Moriya et al., 2017). Erroneous picks were removed after a round of 2D classification with ISAC. After removing bad particles, further segments were removed to ensure that each filament consists of at least 5 members. After 3D refinement, particles were polished using Bayesian polishing routine in Relion (Zivanov et al., 2018) and refined once again within SPHIRE.

### Model building and design

We used the structure of actin in complex with ADP-Pi (PDBID 6FHL) as starting model for the filament, and built a *de novo* model of the Lifeact peptide using Rosetta’s fragment-based approach (Wang et al., 2015). Since one residue of Lifeact is missing from the density, we threaded the sequence in all possible registers – including the reverse orientations – and minimized them in Rosetta. The solution starting from M1 with the N-terminus pointing towards the pointed end was clearly better than all the others (Fig. S2). A set of restraints for phalloidin were built with eLBOW (Moriarty et al., 2009) and the toxin was manually fit into the density with Coot (Emsley et al., 2010). The model was further refined using iterative rounds of Rosetta’s fragment-based iterative refinement (DiMaio et al., 2015), and manual building with Coot and ISOLDE (Croll, 2018; Emsley et al., 2010). The model was finally refined within Phenix (Liebschner et al., 2019) to fit B-factors and correct the remaining geometry errors.

We used RosettaScripts (Fleishman et al., 2011) to design a version of Lifeact with improved affinity. The input protocol and starting files are available upon request. The figures were made using UCSF Chimera (Pettersen et al., 2004).

### Yeast confocal microscopy

Yeast cells bearing plasmids that encode Lifeact-mCherry variants under the native ABP140 promoter and terminator, were grown overnight on a liquid synthetic defined medium (Yeast nitrogen base, Difco) supplemented with glucose. On the following day, the cultures were diluted to OD_600_ 0.5 using fresh media, incubated for 2-3 h at 30 °C and centrifuged at 6,000 g. The cell pellet was washed twice with PBS buffer, fixed by 4% formaldehyde for 20 minutes at room temperature, washed once with PBS buffer and stained with the ActinGreen 488 probe (Invitrogen). After 30 minutes incubation with the probe at room temperature, the cells were washed twice in PBS and applied to concanavalin A-coated glass bottom Petri dishes. Image acquisition was performed with a Zeiss LSM 800 confocal laser scanning microscope, equipped with 63X 1.4 DIC III M27 oil-immersion objective and two lasers with wavelengths of 488 and 561 nm. To avoid experimental bias, we measured the weighted colocalization coefficient of phalloidin with Lifeact-mCherry in 15 cells of each strain using ZEN software.

### Cosedimentation assays

F-actin was prepared as follows. An aliquot of freshly thawed G-actin was centrifuged at 150,000 g using a TLA-55 rotor for 30 minutes at 4 °C to remove possible aggregates. Then, actin was polymerized by incubation in F-buffer (120 mM KCl, 20 mM Tris pH 8, 2 mM MgCl2, 1 mM DTT and 1 mM ATP) for 2 h at room temperature or overnight at 4 °C. For the cosedimentation experiments with myosin, actin was polymerized in the presence of a twofold molar excess of phalloidin. After polymerization, actin filaments were pelleted using a TLA-55 rotor at 150,000 g for 30 minutes and resuspended in the following buffers: F-buffer was used for cosedimentations with tropomyosin or Lifeact-MBP fusion proteins; 20 mM HEPES pH 6.5, 50 mM KCl, 2 mM MgCl_2_ was used for cosedimentations with cofilin; 120 mM KCl, 20 mM Tris pH 8, 2 mM MgCl_2_, 1 mM DTT was used for cosedimentations with myosin.

Cosedimentation assays were performed in 20 μl volumes by first incubating F-actin with the specified proteins for 5 minutes at room temperature, then centrifuging at 150,000 g using the TLA-55 or TLA120.1 rotor for 30 minutes at 4 °C. For the competition assays, Lifeact peptide was added to the mixture at the specified concentrations. After centrifugation, aliquots of the supernatant and resuspended pellet fractions were separated by SDS-PAGE using 4-15% gradient TGX gels (Bio-Rad) and analyzed by densitometry using Image Lab software (Bio-Rad).

### ADP-ribosylation of actin by TccC3HVR

8 μl mixtures of 1 μg (2.4 μM) actin and Lifeact peptide at specified concentrations were pre-incubated for 5 minutes at room temperature in TccC3 buffer (1 mM NAD, 20 mM Tris pH 8, 150 mM NaCl and 1 mM MgCl_2_). The ADP-ribosylation reaction was initiated by addition of 0.02 μg (61 pM) of TccC3HVR into the mixture. After 10 minutes of incubation at 37 °C, the reaction was stopped by adding Laemmli sample buffer and heating the sample at 95 °C for 5 minutes. Components of the mixture were separated by SDS-PAGE, blotted onto a PVDF membrane using a Trans-Blot Turbo transfer system (Bio-Rad) and visualized using a combination of anti-mono-ADP-ribose binding reagent (Merck) and anti-rabbit-HRP antibody (Bio-Rad). The level of actin ADP-ribosylation was quantified by densitometry using Image Lab software (Bio-Rad).

### Confocal microscopy of mammalian cells and the *in vivo* intoxication competition assay

0.05 x 10^6^ HEK 293T cells were seeded in 35 mm glass-bottom, poly-L-lysine coated Petri dishes in 2 mL DMEM/F12 + 10% FBS media and grown for 24 h in a 5% CO_2_ atmosphere at 37 °C. The cells were then transfected with mCherry fusions of actin or Lifeact variants using the FuGENE transfection reagent (Promega). The cells were grown for a further 24 h and transferred to an LSM 800 microscope (Zeiss) equipped with a C-Apochromat 40x/1.2 W objective and maintained in a 5% CO_2_ atmosphere at 37 °C. Images were acquired using the Airyscan detector and a 561 nm laser wavelength for excitation. After taking the first image (0 h), 300 pM PTC3 was added (pre-formed by mixing purified TcdA1 and TcdB2-TccC3 in a 5:1 molar ratio). Images were taken at 10-minute intervals for 5 h.

The images were processed in Fiji (Schindelin et al., 2012). The cells were selected by applying a threshold on the red fluorescence channel, then their areas were measured, normalized to the original cell area at the 0 h time point, and the resulting % change in cell area that occurred during the experiment was plotted.

Three independent experiments for each construct were performed. The cells were not tested for *Mycoplasma* contamination.

## Figure legends

**Figure S1.**
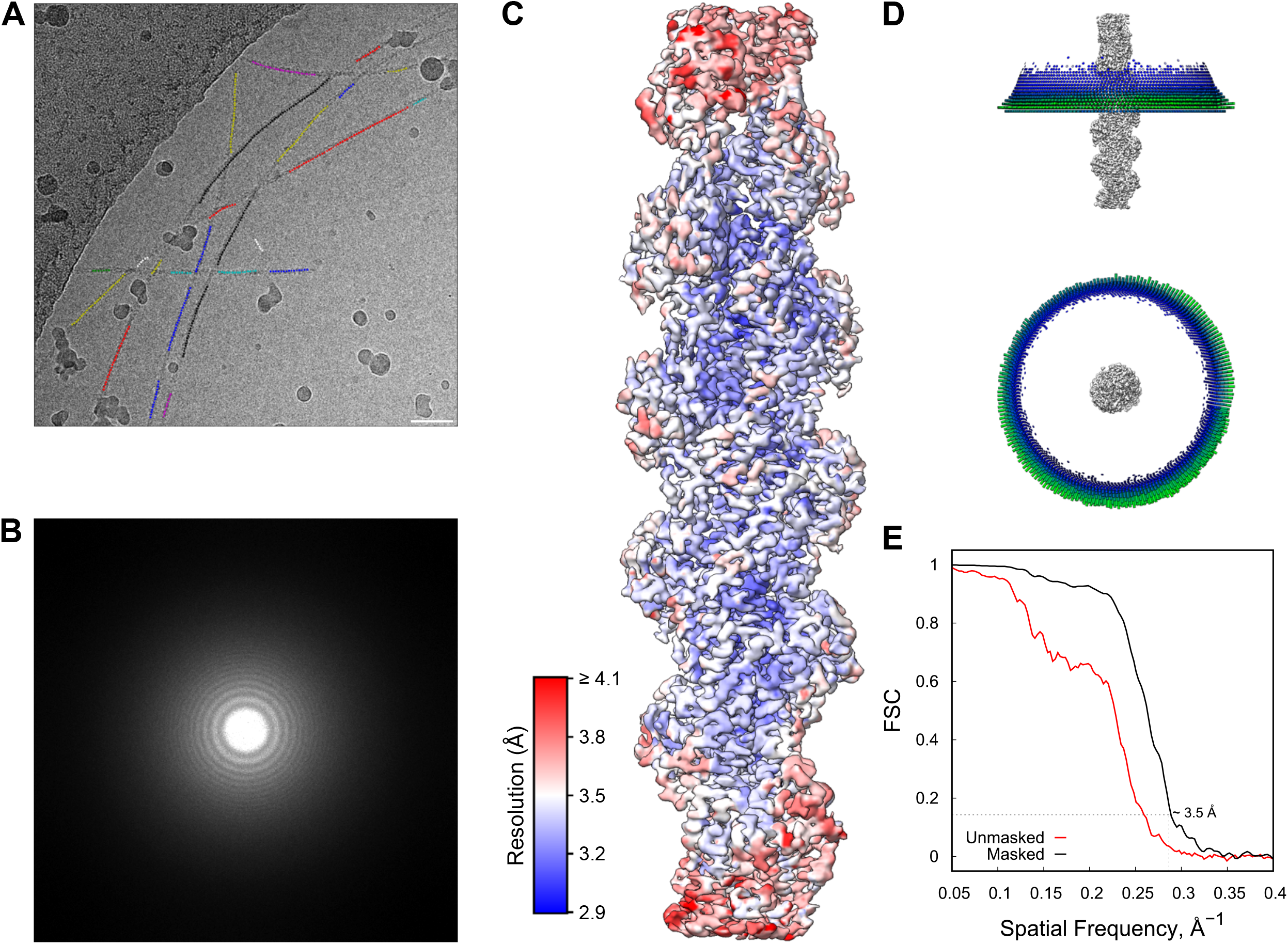
Overview of the cryo-EM data. **A, B:** Example micrograph (**A**) and its power spectrum (**B)** at ~ −1.5 μm defocus. Filaments selected automatically by crYOLO are highlighted as differently colored dots. Scale bar 10 μm. **C**: Density map of the Lifeact-F-actin complex colored according to the local resolution. **D**: Orientation distribution of the particles used in the final refinement round. **E:** Fourier shell correlation (FSC) for the masked and unmasked final reconstructions. The FSC was calculated in the central 120 Å area of the map.

**Figure S2.**
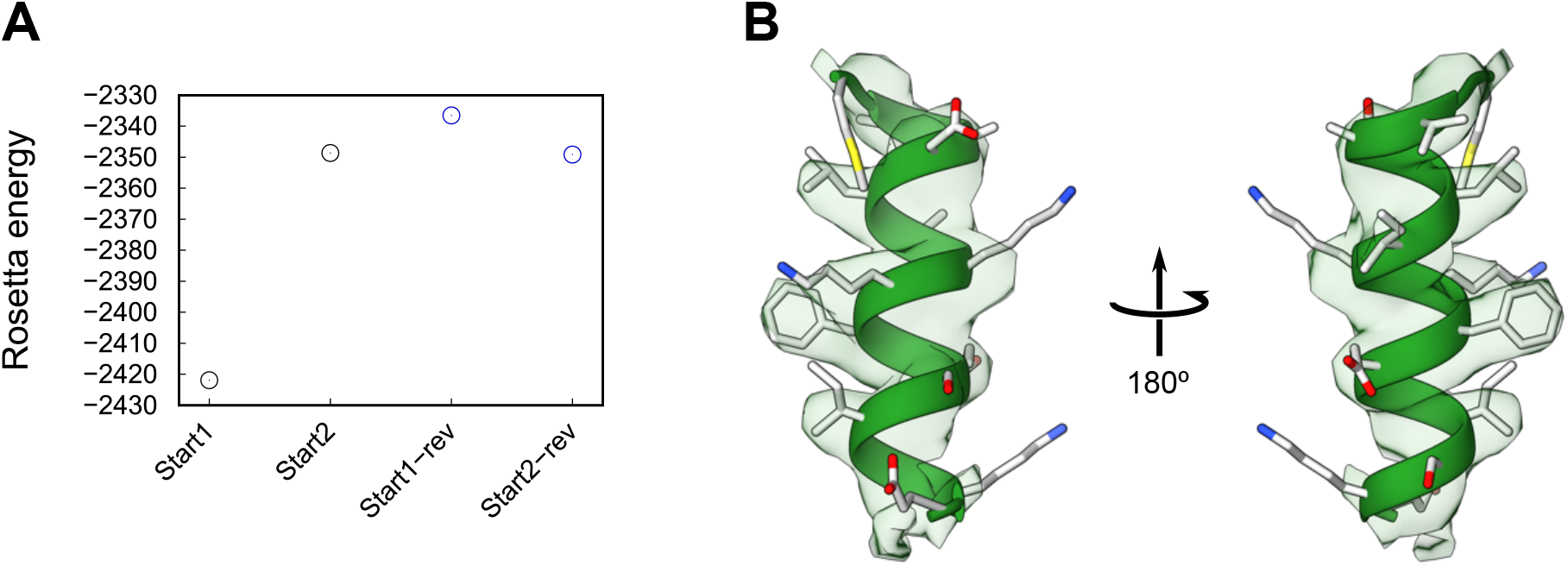
Fit quality of the atomic model of LifeAct. **A:** Minimized energy of models of all possible registers of the LifeAct sequence into the density. Start1 and 2 correspond to the peptides starting at M1 or G2 with their N-termini pointing towards the pointed end of the filament. For Start1-rev and Start2-rev the N-termini points towards the barbed end. The Rosetta energy values show a clear preferred solution. **B:** Density fit of the final model corresponding to the energy minimum seen in (**A**).

**Figure S3.**
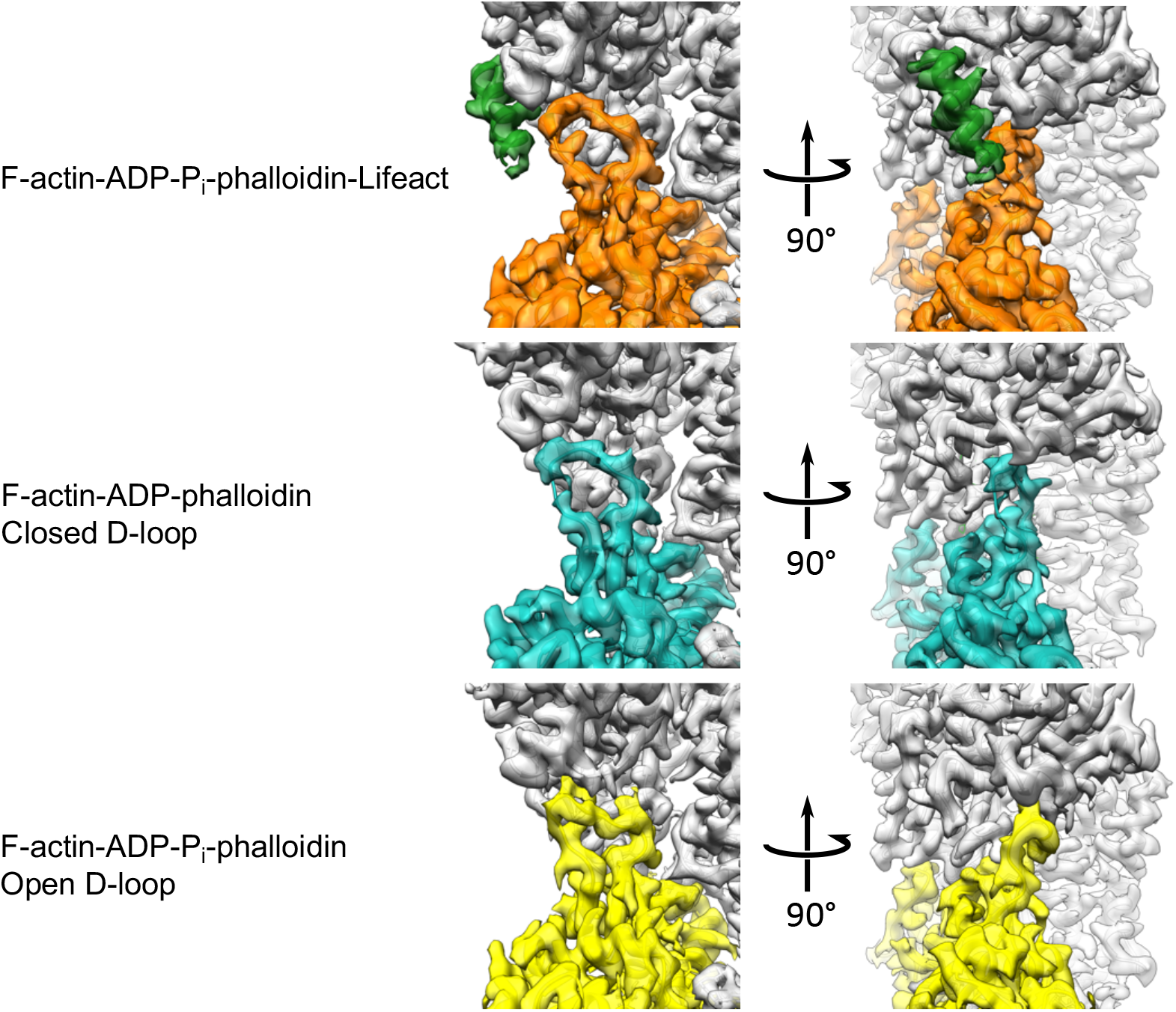
Comparison of D-loop conformations. The density maps and corresponding atomic models of Lifeact-F-actin-ADP-Pi-phalloidin in comparison to those of the open D-loop state in F-actin-ADP-Pi-phalloidin (PDB 6T1Y, EMDB 10363) (Pospich et al., 2019) and closed D-loop state in F-actin-ADP-phalloidin (PDB 6T20, EMDB 10364) (Pospich et al., 2019).

**Movie S1**. **Stabilization of the D-loop in its closed state upon Lifeact binding. A:** The density map of F-actin-ADP-Pi-phalloidin (EMDB 10363, gray) is superimposed with the corresponding atomic model (PDB 6T20) (Pospich et al., 2019). **B**: Close-up view of the D-loop and the C-terminal interface of F-actin-ADP-Pi-phalloidin complex. Possible position of Lifeact on F-actin-ADP-Pi-phalloidin shown in transparent green. Note that the D-loop is in its open state. **C**: Close-up view on the interface between Lifeact and F-actin-ADP-Pi-phalloidin complex. I13 of Lifeact interacts with M47 of F-actin, stabilizing the closed D-loop conformation in F-actin. **D**: Overview of the complete Lifeact-F-actin-ADP-Pi-phalloidin map and the corresponding atomic model.

## Author contribution

A.B. prepared cryo-EM specimens and collected data. F.M. processed cryo-EM data, built the atomic model and design. A.B. performed *in vitro* competition and activity assays, and *in vivo* experiments in yeast. O.S. performed and analyzed intoxication assays in mammalian cells. A.B, F.M. and O.S. prepared figures and video. A.B., F.M. and O.S. wrote the original draft of the manuscript. S.R. supervised the project, reviewed and edited the manuscript. All authors reviewed the results and commented on the manuscript.

## Acknowledgements

We thank S. Shydlovskyi for assistance with data collection, O. Hofnagel and D. Prumbaum for maintaining the EM facility, S. Bergbrede and M. Hülseweh for the excellent technical assistance, W. Linke and A. Unger (Ruhr-Universität Bochum, Germany) for providing us with muscle acetone powder, S. Rospert (University of Freiburg, Germany) for providing us anti-RPS9 serum, and S. Pospich for fruitful discussions. This work has been funded by the Max Planck Society (S.R.). A.B. is a fellow of the Humboldt foundation.

## Supplementary materials and methods

**Table S1.**
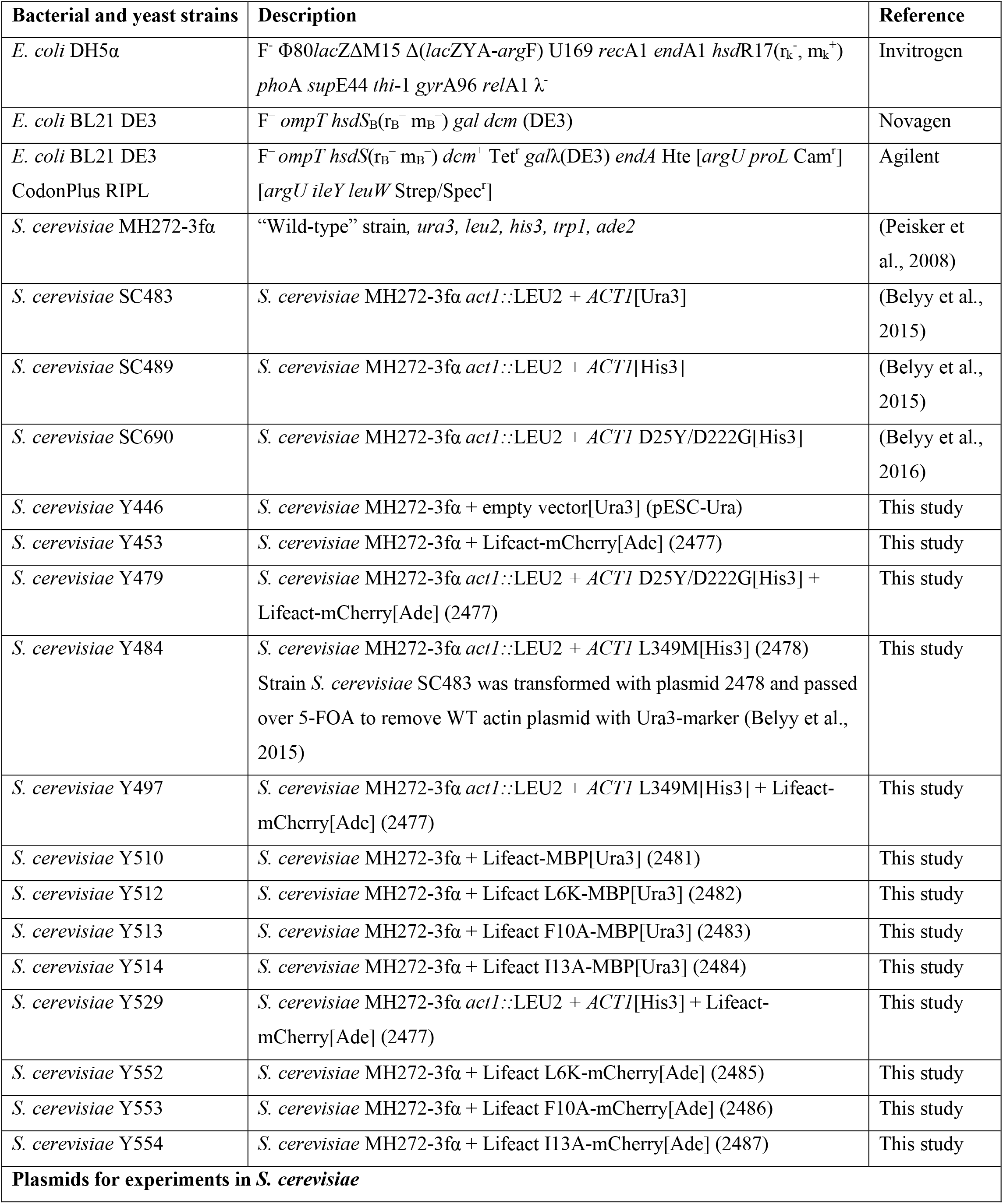
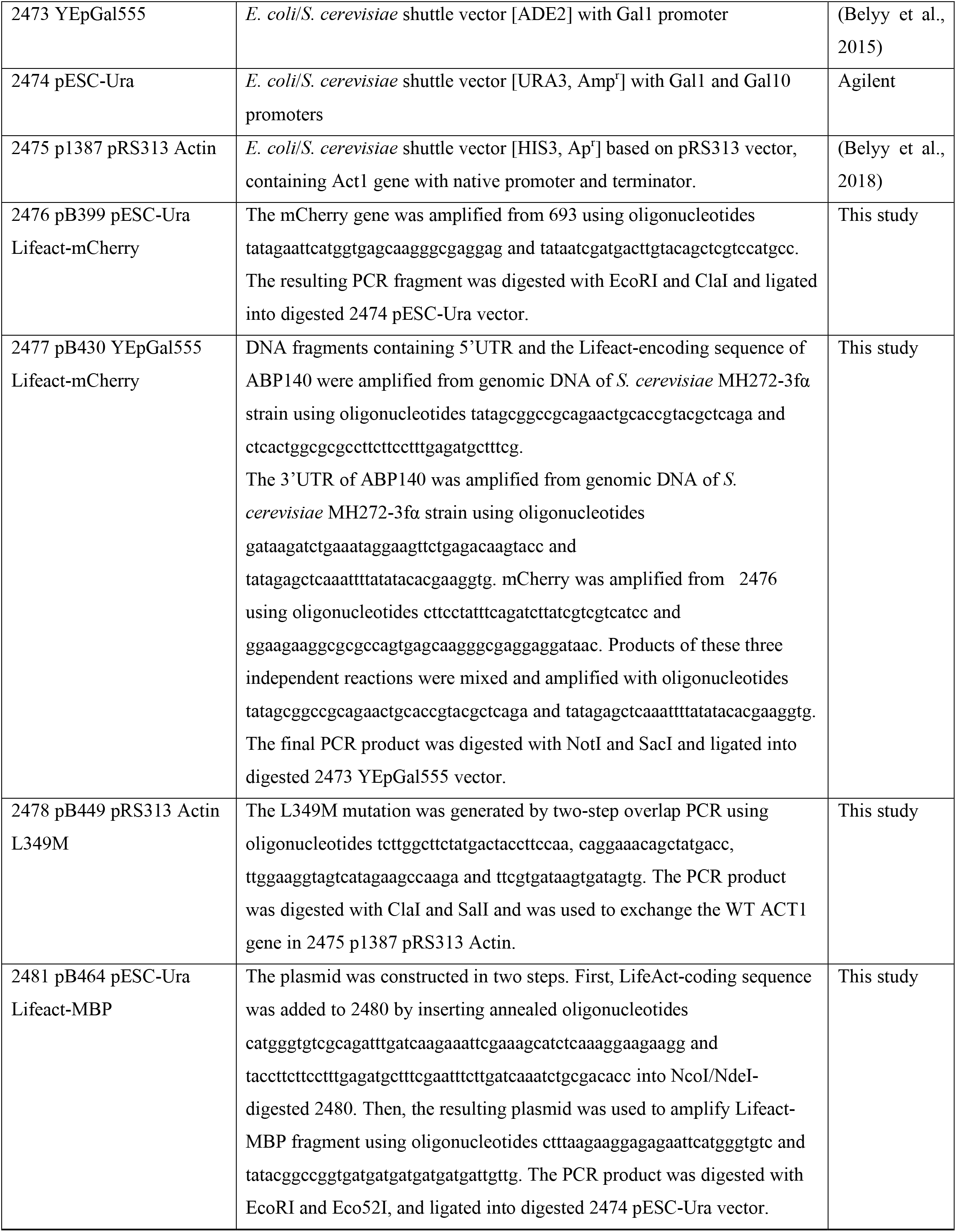
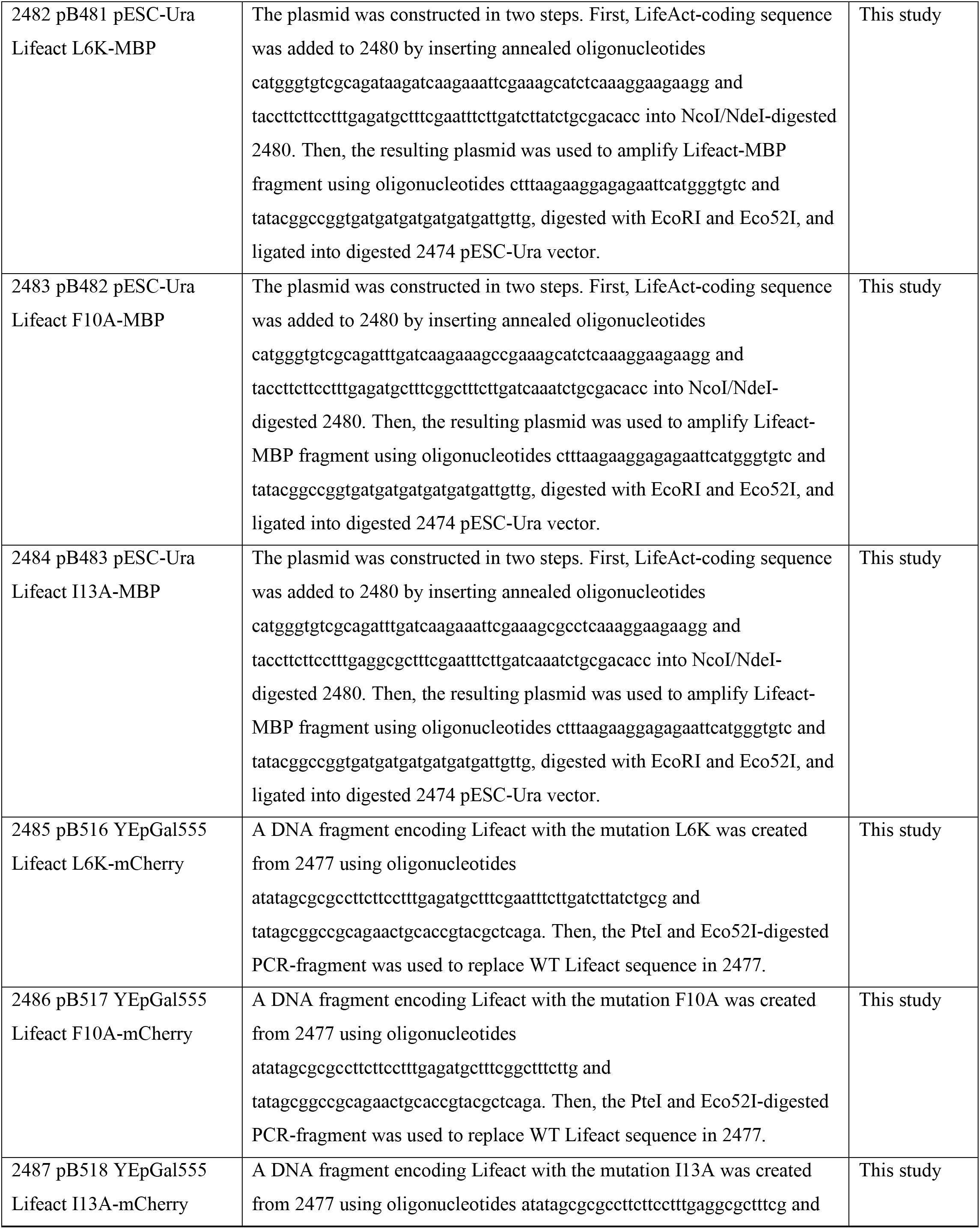
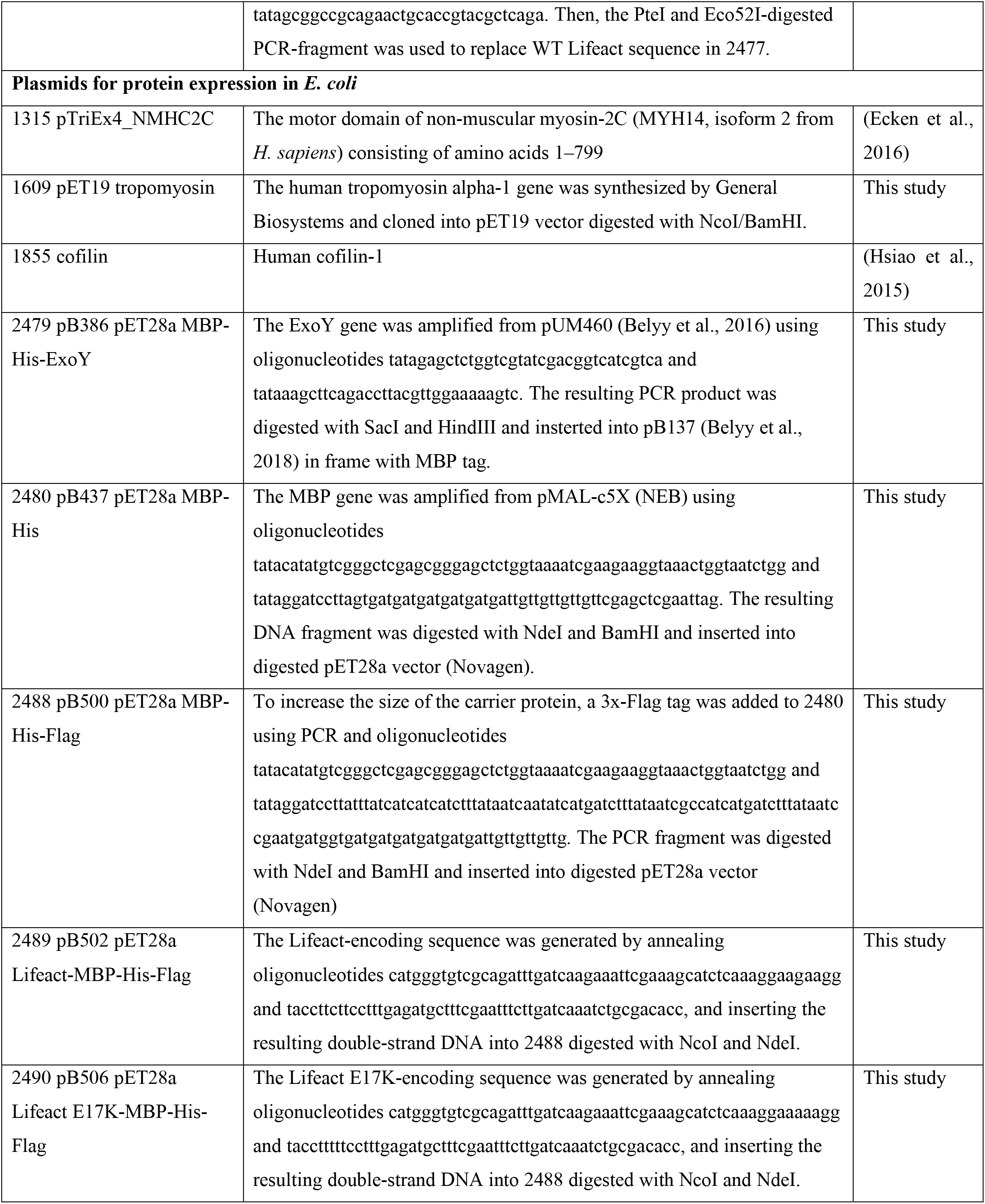
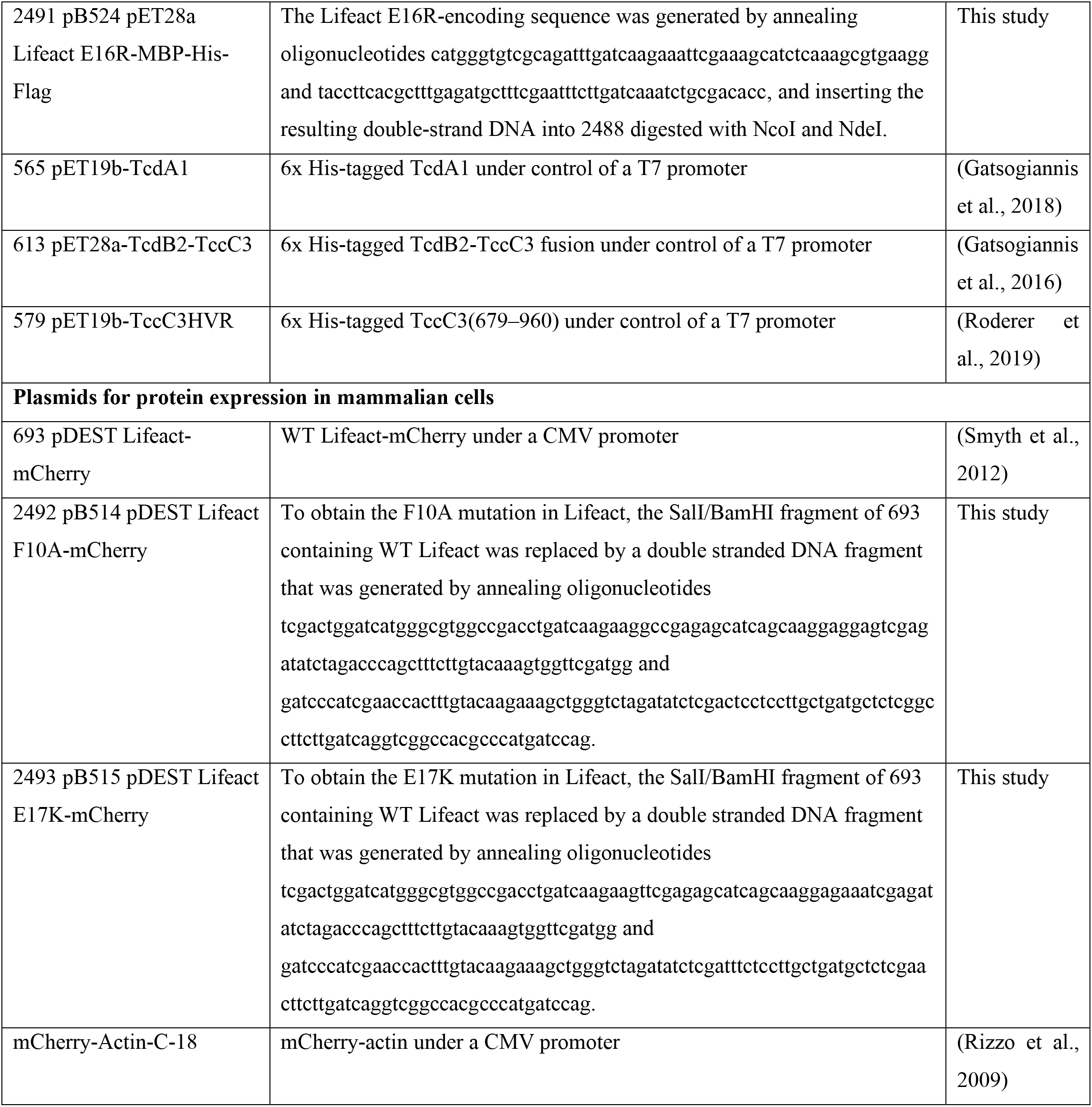
List of primers, strains and plasmids used in this study.

